# Real time genetic compensation operationally defines the dynamic demands of feedback control

**DOI:** 10.1101/244020

**Authors:** Patrick Harrigan, Hiten D. Madhani, Hana El-Samad

## Abstract

Biological signaling networks use feedback control to dynamically adjust their operation in real time. Traditional static genetic methods such as gene knockouts or rescue experiments often can identify the existence of feedback interactions, yet are unable to determine what feedback dynamics are required. Here, we implement a new strategy, closed loop optogenetic compensation (CLOC), to address this problem. Using a custom-built hardware and software infrastructure, CLOC monitors in real time the output of a pathway deleted for a feedback regulator. A minimal model uses these measurements to calculate and deliver—on the fly—an optogenetically-enabled transcriptional input designed to compensate for the effects of the feedback deletion. Application of CLOC to the yeast pheromone response pathway revealed surprisingly distinct dynamic requirements for three well-studied feedback regulators. CLOC, a marriage of control theory and traditional genetics, presents a broadly applicable methodology for defining the dynamic function of biological feedback regulators.

## INTRODUCTION

The classical genetic compensation experiment tests whether phenotypes produced by a mutation can be rescued by the wild-type allele (Collins et al., 2006). This experiment, however, does not reveal whether the *time-dependent* regulation of the corresponding gene or gene product is important for maintaining the wild-type phenotype. Such knowledge is particularly germane to feedback regulation in signaling and transcriptional networks where the dynamic interplay between pathway activity and feedback activation enables cells to monitor and adjust the activity of upstream components in real time. Feedback regulation is a widespread cellular strategy that underlies diverse cellular behaviors including perfect adaptation (Leibler et al., 1999), multi-stability (Xiong and Ferrell, 2003) and oscillations (Tsai et al., 2008). In these contexts, feedback plays a dominant qualitative role that can be assessed by inference since its loss usually induces visible disruption of a phenotype—for example loss of oscillations (Lahav et al., 2004) or perfect adaptation (Muzzey et al., 2009). However, the majority of feedbacks loops seem to play quantitative roles, fine tuning the operation of cellular pathways to demands and fluctuations in the environment (Nevozhay et al., 2009). Such regulation is challenging to understand quantitatively because the input-output relationship of many interconnected signaling components must be measured and modeled mathematically (Garmendia-Torres, Goldbeter and Jacquet, 2007; Howell et al., 2012). Mathematical predictions need then to be iteratively tested in painstaking fashion (Harreman et al., 2004; Cirit, Wang and Haugh, 2010; Dixit et al., 2014).

We describe here a general method to study cellular feedback regulation that employs and updates the conceptual framework of genetic compensation experiments (**Figure 1A**). In this approach, we define a quantitative, dynamic phenotype associated with the loss of a pathway-controlled regulator: the time-dependent difference between the output of the wild type and mutant pathways. We use this information and a minimal mathematical model to compute continuously, for every time point at which the output is measured, how much of the missing regulator needs to be added back into the cell in order to rescue this phenotype and restore wild-type output dynamics. We then inject the computed level of the regulator into the mutant cell using an optogenetic construct that controls its transcription. In this closed loop optogenetic compensation (CLOC) approach, both fully and partially compensatory inputs produce focused hypotheses about the temporal demands on the feedback regulator. Below we describe how we have applied CLOC to three well-studied negative regulators of the yeast pheromone response MAPK pathway, a dual-specificity MAPK phosphatase Msg5, a DEP-RGS protein Sst2, and the Gα protein Gpa1. Our studies reveal distinct dynamic requirements for each of these pheromone-induced regulatory factors and reveals the power of elucidating the time scales of gene function, which cannot be fully interrogated by compensation with static genetic alleles.

**Figure 1.**
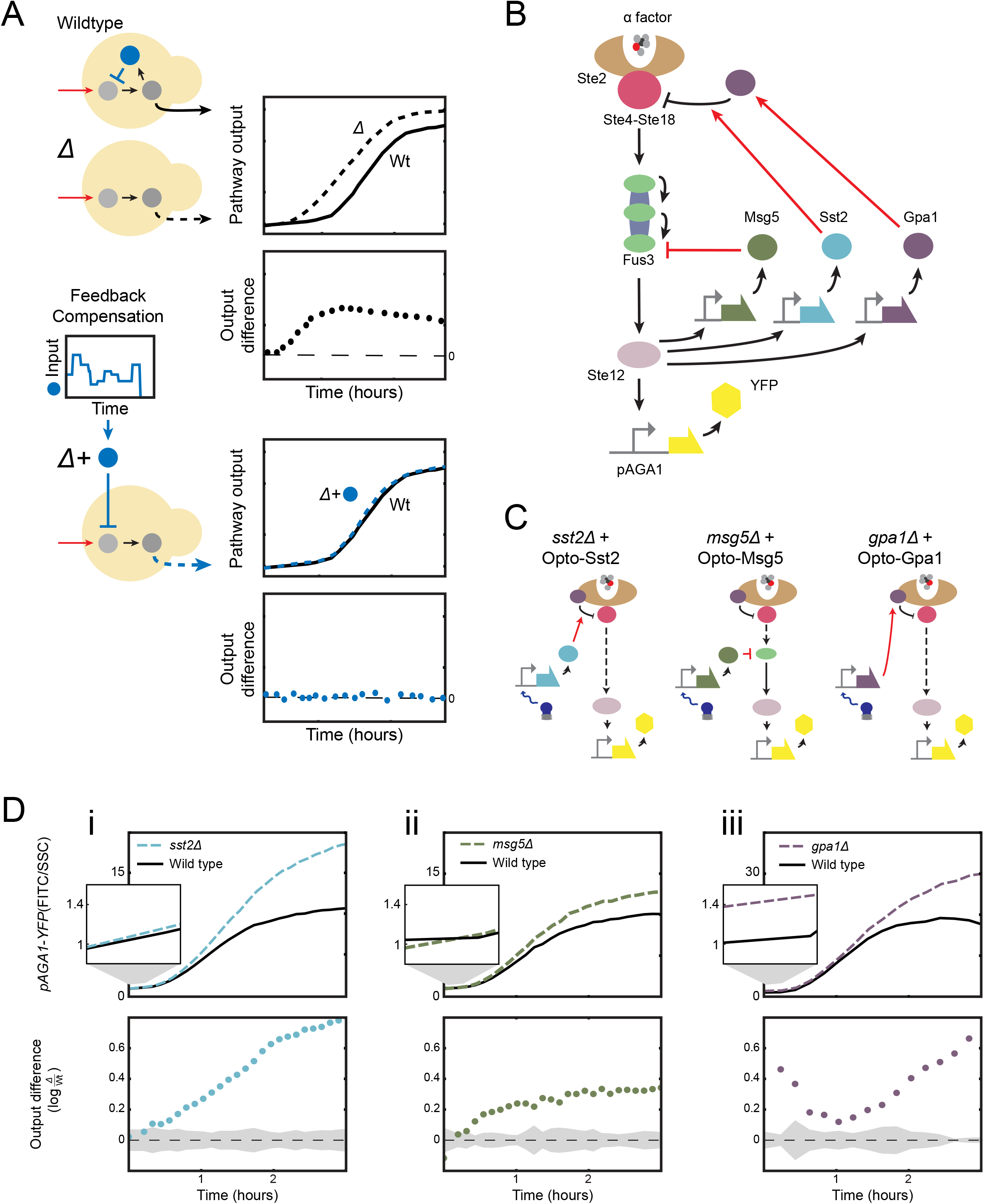
Mutants of *SST2*, *MSG5*, and *GPA1*, three pheromone-induced negative regulators of the yeast mating pathway have different signaling dynamics phenotypes. (a) Illustration of the concept of time dependent genetic compensation. Deletion of a feedback regulator causes a difference in the pathway output of the mutant as compared to the wildtype. This difference is time-varying. As a result, rescue of this quantitative dynamic phenotype requires the time-varying exogenous expression of the missing gene. (b) Schematic of the wild type pheromone response pathway. (c) The different mutants for pheromone responsive regulators (*SST2*, *MSG5* and *GPA1*). Rescue of these mutations will be carried out using an optogenetic construct. (d) The α-factor response for *sst2*∆ (i), *msg5*∆ (ii), and *gpa1*∆ (iii) compared to the wild type response. Top panels: Fold change *pAGA1-YFP* expression as a function of time for the wild type and feedback deficient strain after pathway induction with 0.5nM α-factor. Bottom Panels: Difference of *pAGA1-YFP* expression in the wild type and feedback deficient strains. Zero indicates no difference. Shaded region represents the standard deviation when comparing three biological repeats of wild type signaling. This region will help us define successful compensation to wild type given experimental error and biological variability.

## RESULTS

### *In silico* feedback control of yeast mating pathway dynamics by automated optogenetic expression of negative regulators

The yeast mating pheromone response pathway is a model signal transduction network(Bardwell, 2005) where extensive work mapping pathway components and interactions has identified numerous feedback regulators (Kaffman, Rank and O’Shea, 1998; Ren et al., 2000; Gruhler et al., 2005). In this pathway, the binding of mating pheromone to the G-protein-coupled receptor Ste2 (**a**-cells) or Ste3 (α-cells), is transduced by a MAP kinase cascade into the activation of the transcription factor Ste12. Ste12 drives the expression of genes required for mating as well as genes encoding for pathway components that regulate upstream signaling. We investigate three such negative regulators whose transcription is stimulated by pheromone: Sst2 (Dietzel and Kurjan, 1987), Msg5 (Doi et al., 1994), and Gpa1 (Nakayama et al., 1988) (**Figure 1B**). Gpa1 is the alpha subunit of the receptor-controlled G protein that inhibits the Gβγ complex (Ste4-Ste18), which, once freed from inhibition, is the key activator of the MAPK cascade. Msg5 is the dual specificity phosphatase that dephosphorylates and inactivates the MAPK Fus3. Sst2 is a DEP-RGS protein that enhances GTP hydrolysis by Gpa1, terminating signaling and promoting receptor down-regulation. While these three negative regulators are well-studied, how their *temporal regulation* by the MAPK pathway shapes signaling dynamics is not well-understood.

To study the dynamic requirements of these pheromone controlled pathway regulators, we constructed feedback deficient mutants (*sst2*∆, *msg5*∆, *gpa1*∆) in which the endogenous Ste12 inducible copy of each gene is knocked out and replaced with a corresponding optogenetically inducible copy (**Figure 1C**). This effectively replaces the native Ste12-activated expression of the feedback regulators with expression that can be experimentally activated with light. Light-based inputs, which can be easily varied in time and magnitude, are particularly well suited for fine, dynamic control of gene expression. We used a modified version of a previously reported cryptochrome based optogenetic expression system(Kennedy et al., 2010) in which the cytochrome protein Cry2phr is fused to the Gal4 DNA binding domain and its interacting partner Cib1 is fused to the activation domain of Rtg3 (Rothermel, Thornton and Butow, 1997). Upon absorption of a photon of blue light, Cry2phr-Gal4DBD undergoes a conformational change that enables it to bind Cib1-Rtg3AD and drive expression from the p*GAL1* promoter of either Sst2 (Opto-Sste2), Msg5 (Opto-Msg5), or Gpa1 (Opto-Gpa1). The Cry2phr-Gal4DBD optogenetic construct was effective at expressing these proteins, and we observed robust and dose dependent repression of pathway output upon exposure to blue light, as measured by a pheromone-responsive *pAGA1-YFP* reporter (Mccullagh et al., 2010) (**Figure S1-S3**)

To assess the impact of removing SST2, MSG5 or GPA1 on mating pathway signaling dynamics, we used α-factor to induce each feedback deficient mutant in the dark (with no induction of Opto-Msg5, Opto-Sst2, or Opto-Gpa1) and compared its pathway output to that of the wild type (**Figure 1D**). In *sst2*∆ cells, the difference in pathway output relative to wild-type increases steadily over time while in *msg5*∆ cells this difference remains roughly constant for the duration of the experiment after an initial rise. Unlike the other mutants, *gpa1*∆ cells display a quantitative and reproducible increase in basal signaling prior to pheromone addition. This initial difference in pathway activity decreases during early induction and then increases as *gpa1*∆ reaches a higher maximum pathway activity than wild type.

One strategy for restoring wild type pathway dynamics in the feedback deficient mutants would be to iteratively test predetermined optogenetic inputs until one that restores wild type output is found. This strategy, referred to in engineering as open-loop control, requires a predictive understanding of how pathway output changes as a function of optogenetic input and can therefore be unreliable if this function is affected by day to day variation in pathway behavior (Milias-Argeitis et al., 2011b, 2016; Del Vecchio et al., 2017; Rullan et al., 2017). The alternative ‘closed-loop’ approach would leverage the same strategy that endogenous cellular feedback loops use to produce robust signaling dynamics—monitor the pathway output in real time and correspondingly adjust the activity of the feedback regulators (**Figure 2**). In our case, this entails real-time, *in silico* adjustment of the optogenetic input to change *in vivo* regulator expression and in this fashion, staple the output of the mutant to that of the wild type. Similar closed loop control schemes have recently been used to control intracellular processes in live cells (Milias-Argeitis et al., 2011a; Toettcher et al., 2011; Uhlendorf et al., 2012). Historically, they form the basis of patch clamp studies in neurobiology (Hodgkin and Huxley, 1952) where a controller-supplied current that is updated in real-time to compensate for neuron voltage changes serves as a proxy for measuring endogenous neuron currents. Likewise, we hypothesize that the dynamic optogenetic input required for time-dependent genetic compensation of a feedback regulator serves as a proxy for the deleted endogenous feedback, allowing for quantitative characterization of the feedback regulator’s temporal requirements in the context of native pathway signaling.

**Figure 2.**
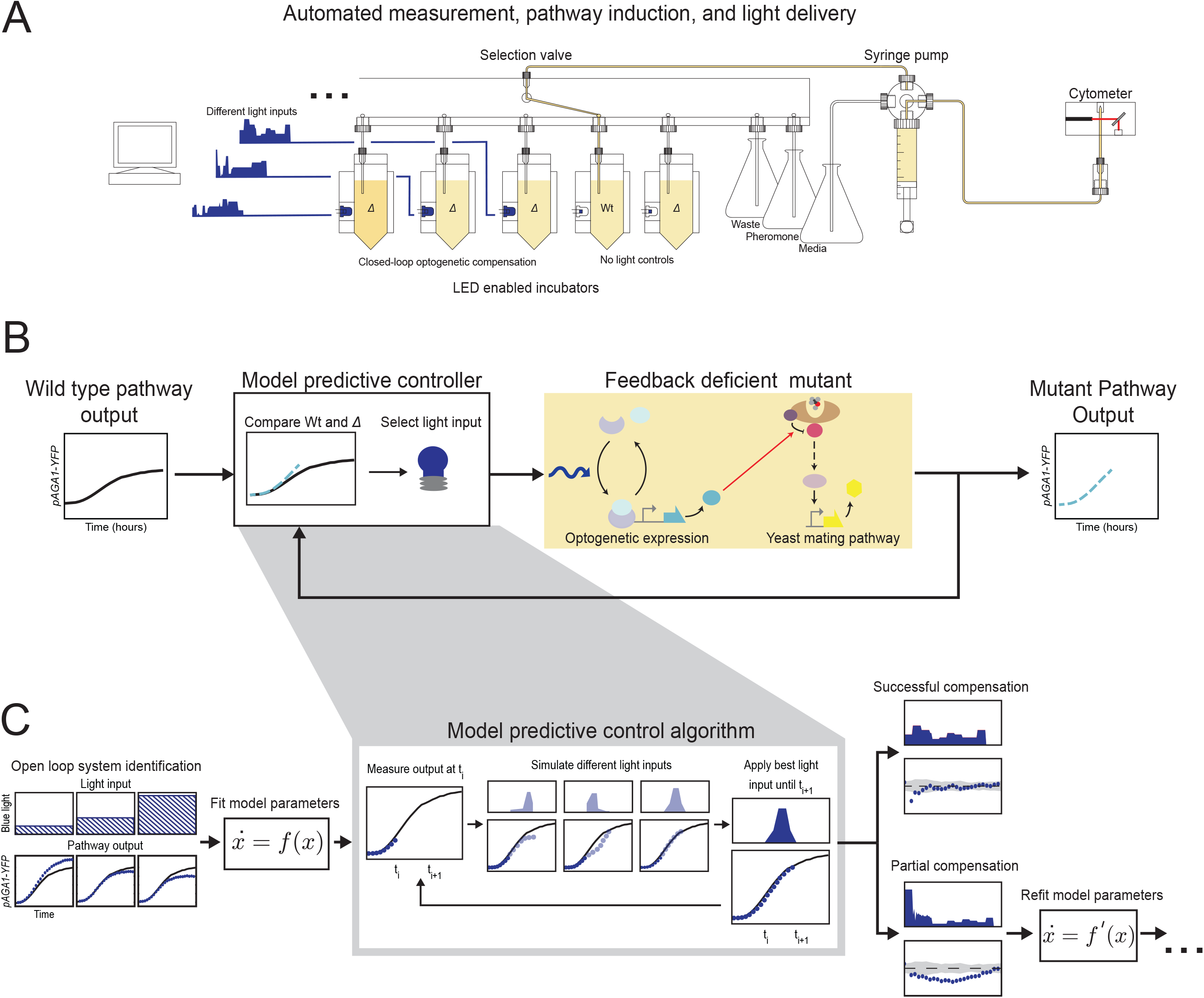
Synthetic optogenetic compensation control (CLOC). (a) Schematic showing the LED-enabled yeast incubation array and liquid handling device used for real time cytometry measurement and light delivery. (b) Block diagram showing the design of the closed control strategy used to implement CLOC. An *in silico* computational controller continuously compares desired pathway output to mutant pathway output and determines the appropriate optogenetic expression of a pathway regulator to carry out the genetic compensation of the mutant output. (c) Schematic of model identification and model predictive control. The output of the mutant pathway in response to constant light input is used to parameterize an initial model. Model predictive control determines appropriate light inputs to carry out the dynamic compensation in an initial set of CLOC experiments. If compensation is not satisfactory, model identification is repeated with all generated data, including suboptimal CLOC trials. Eventually, CLOC trials converge on a satisfactory dynamic compensation strategy. (See Table S4)

To accomplish closed-loop control, we designed and fabricated an automated yeast culturing, light delivery, and cytometric measurement device (**Figure 2A**). The device consists of 8 temperature-controlled and aerated yeast incubators with individually addressable and integrated blue light-emitting diodes (LED) for light delivery. A selection valve and syringe pump were used to automate the addition of fresh media and α-factor to the cultures as well as the injection of culture into a flow cytometer to measure pathway activity. The device was controlled via customized code that allows the user to set experimental parameters including measurement frequency and the timing of α-factor introduction (**Supplemental Information**).

The device uses a particular closed loop control strategy, model predictive control (MPC) (Borrelli, Bemporad and Morari, 2017), to determine the appropriate optogenetic input for CLOC from real-time output measurements of the mating pathway (**Figure 2B**). Although MPC requires the construction of a rudimentary model of the system, it has the advantage over system-agnostic control schemes such as the classical proportional-integral-derivative method in that it is better suited for systems with significant delays such as those inherent to transcription and translation. We chose a two-state ordinary differential equation (ODE) model that yielded a simplified description of how *pAGA1-YFP* expression responds to both mating pheromone (α-factor) and light in the feedback deficient mutants (**Supplemental Information**). The two variables described in the model represent the dynamics of light induced expression of the feedback regulator and the measured fluorescent output in response to this regulator. This model is a parsimonious and generic description, and we chose it with the explicit intent of developing a generalized approach *without* the need for detailed mechanistic models. During a typical CLOC experiment, we induce signaling using α-factor and measure the *pAGA1-YFP* output via flow cytometry. The MPC controller uses this measurement to estimate the states of the ODE model and then selects a light input that the model predicts minimizes the future difference between the activity of the mutant pathway and the desired wild type pathway (**Figure 2C**). This light input is then applied to the mutants for 15 minutes, at which point the operation is repeated. The states of the model are re-estimated based on the current measurements of *pAGA1-YFP* and a new light input is selected and administered.

Using our automated closed-loop control device, we performed time dependent genetic compensation of *sst2*∆, *msg5*∆, and *gpa1*∆ with Opto-Sst2, Opto-Msg5, and Opto-Gpa1 respectively. In each case, we initially tested constant-light induction of the negative regulator in open loop—that is without changing light input as a function of *pAGA1-YFP* measurements—for a range of light intensities. The open loop response of the mutants was then used for the initial parameterization of the controller ODE model (**Figures S1-S3**). In cases where this parameterization was insufficient for obtaining satisfactory CLOC, we adopted a procedure in which we iteratively refit the model parameters using data from CLOC experiments that only achieved partial compensation (**Figures S2-S3**). This is akin to closed-loop system identification (Ljung, 1999), a procedure used to ensure that a low resolution empirical model still captures the properties of a given system (such as important time-scales) that are required for efficient closed loop control. Examination of these iterative trials that achieved either satisfactory or partial compensation revealed striking differences for the dynamic requirements of the three regulators.

### Constant Opto-Sst2 induction rescues wild-type dynamics

We first attempted to rescue wild-type signaling dynamics in an *sst2*∆ mutant using constant open loop induction of Opto-Sst2. Given the often-assumed canonical role of *SST2* as a feedback regulator (Chasse et al., 2006), we were surprised to find that constant induction of Opto-Sst2 at an appropriate light intensity was sufficient to rescue wild type signaling dynamics in *sst2*∆ (**Figure 3A**). Constant induction above or below this level achieved partial compensation, initially driving signaling to wild type levels before eventually resulting in pathway over attenuation or over activation compared to wild type (**Figure 3A trials 1–5**). These data strongly indicate that constant pheromone-induced expression of *SST2* suffices to program the wild-type response to mating pheromone and that this response does not require a dynamic transcriptional feedback mechanism. This is further corroborated by the light inputs used in CLOC, which resemble fluctuations about a constant light intensity (**Figure 3A trial 6**). Constant induction of Opto-Sst2 was similarly sufficient in rescuing wild type dynamics for experiments performed at a higher concentration of α-factor (**Figure S4A**).

**Figure 3.**
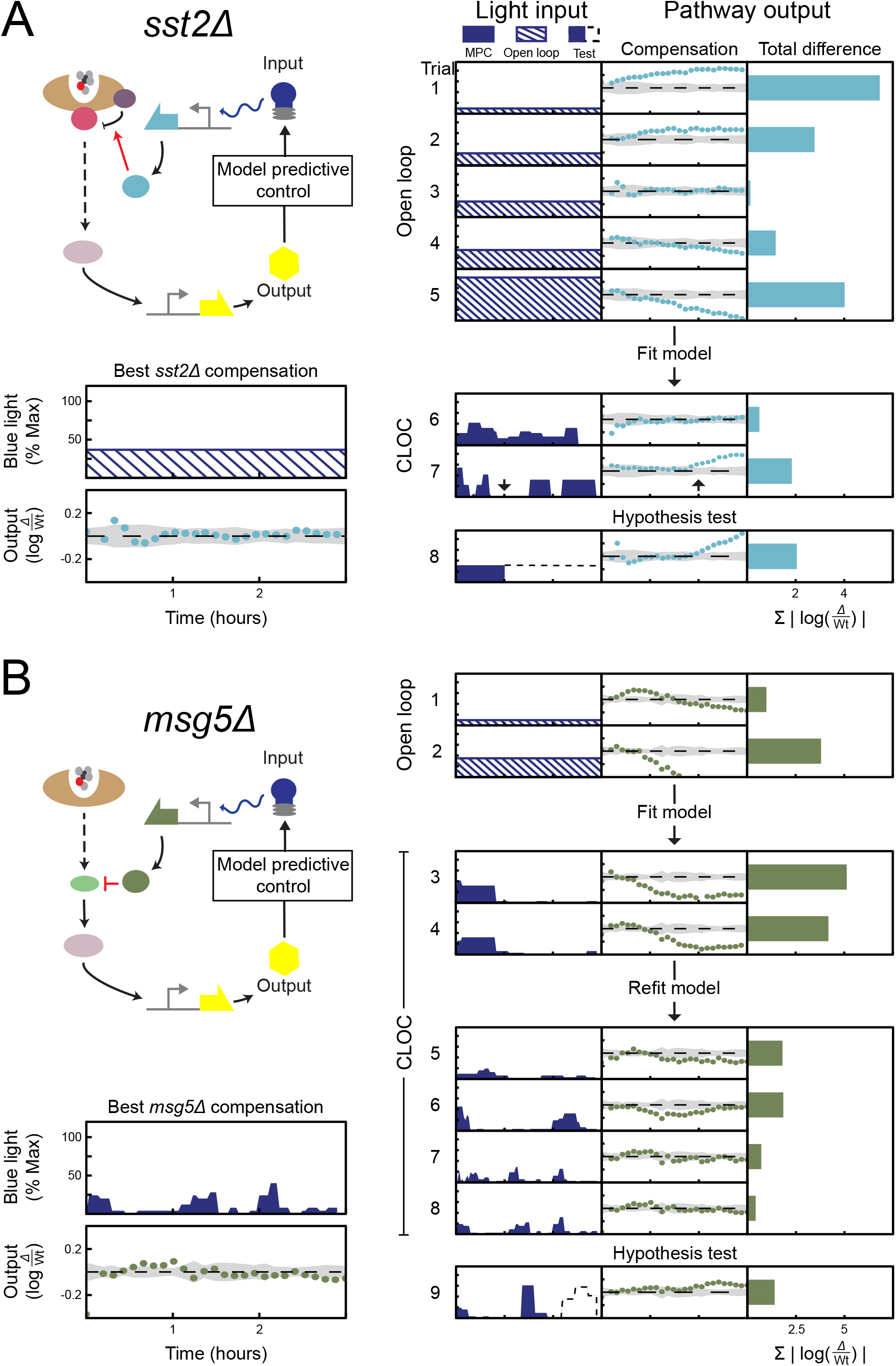
CLOC experiments indicate that there are no dynamic requirements for *SST2*. **(a). By contrast, dynamic compensation of *MSG5* is necessary to rescue wild type pathway output (b)..** For all trials, mutant strains are induced with 0.5 nM α-factor at time 0 hours and *pAGA1-YFP* is then measured for 3 hours. Compensation of mutants is carried out through optogenetic expression of the deleted regulator. Results of different trials are shown as rows in tables with three columns. The first column shows the light input as a function of time, the second column shows the resulting difference in signaling output between compensated mutant and wild type as a function of time, and the third column shows the sum of the differences shown in the second column. Light inputs were given in either open loop (input is dashed), closed loop (input is solid color). For hypothesis testing, input prescribed was implemented (input in blue solid color) until some point in time when the control was purposefully shut off (input in dashed black line). The optogenetic input resulting in the best compensation along with its compensation error is shown on the left. This plot also shows the numerical values of the axes. The same scale and values are used for all panels. (See Figure S1, Figure S2, Figure S4, and Table S3)

Further examination of the MPC inputs revealed an interesting trend: even when induction of opto-Sst2 is switched off for a period of time during CLOC, we didn’t observe a corresponding over activation of *sst2*∆ signaling until approximately one hour later. (**Figure 3A trial 7**). To directly test this observed delay, we carried out an experiment in which we expressed Opto-Sst2 at the optimal level for 1 hour and then turned off its induction. We observed that there is indeed a one hour delay before *sst2*∆ output deviates from wild-type levels (**Figure 3A trial 8**). This slow pathway output to shutoff of Opto-Sst2, which could be related to the half-life of Sste2, the responsiveness of the pathway, or both, is consistent with the apparent lack of a temporal requirement for the regulation Sst2.

### Dynamic requirement for Opto-Msg5 to rescue wild-type signaling

In contrast to *SST2*, we could not find any level of constant Opto-Msg5 expression that rescues wild type signaling dynamics in *msg5*∆ (**Figure S2A**). Instead, we observed a tradeoff in which a constant induction of Opto-Msg5 that is strong enough to drive early *msg5*∆ signaling to wild type levels also resulted in over attenuation of the output at later times (**Figure 3B trial 2**). Dialing down the constant light intensity could reduce this over-attenuation but at the cost of initial over-activation (**Figure 3B trial 1**). The tradeoff between the need for sufficient initial attenuation and its incurred subsequent over-attenuation suggested that a dynamic opto-Msg5 input was needed to circumvent this limitation of constant induction.

We then carried out CLOC experiments using a model parameterized from the collected open loop data. In these first experiments, the MPC attempted to prevent the problem of over attenuation by stopping induction of Opto-Msg5 after early *msg5*∆ signaling had been restored to wild type levels (**Figure 3B trials 3–4**). However, this still resulted in poor compensation as pathway signaling was slow to recover from its initial attenuation even in the absence of opto-Msg5 induction. This is likely due to the excess Opto-Msg5 molecules produced during the overzealous initial pulse that persist in the cell and continue to repress pathway output for an extended period of time. This suggested that initial expression of Msg5 is crucial for wild type signaling dynamics and that mistakes of over-production cannot be promptly corrected. Furthermore, it was evident that the model parameters used in CLOC were not sufficiently tuned to account for the important timescales of the signaling process.

Therefore we refitted the model parameters using these initial CLOC trials. With the refitted model, CLOC converged on a strategy of periodic pulsing of Opto-Msg5 expression (**Figure 3B trials 7–8**). This strategy alleviated the shortcomings of previous trials by adjusting the first Opto-Msg5 pulse duration and timing to drive initial pathway output to the level of wild type without excess production of Opto-Msg5. Subsequent Opto-Msg5 pulses were then administered as the impact of the previous pulse of Opto-Msg5 is lost. This interpretation is supported by non-optimal CLOC trials in which the initial pulse of Opto-Msg5 induction was too long (**Figure 3B trial 5**) or strong (**Figure 3B trial 6**) and drove pathway output below wild type. In both cases, the second Opto-Msg5 was not administered. We again observed similar results for CLOC experiments conducted at a higher concentration of α-factor (**Figure S3B**). Taken together, these experiments indicate that dynamic pulses of *MSG5*-mediated feedback enable fine control of pathway dynamics while avoiding over attenuation.

To further test this hypothesis, we conducted CLOC experiments in which the LEDs were purposefully turned off before the MPC could administer the third pulse seen in successful CLOC trials (**Figure 3B trial 9**). Without a third pulse, we consistently observed that pathway activity rose above the wild type level at the end of the experiment.

Interestingly, while different successful CLOC experiments always generated a pattern of dynamic pulsing, the magnitude and timing of these pulses varied quantitatively between experiments, likely as a response to biological variability in the pathway (Milias-Argeitis et al., 2016). Our control scheme is able to account for this variability and adjust quantitatively to it in real-time. This suggests that the endogenous *MSG5* feedback loop might carry out the same highly dynamic function in cells and further supports the hypothesis that dynamic expression of Msg5 in feedback is crucial to maintain wild type signaling demands in the pathway.

### Dynamic requirement for Opto-Gpa1 to rescue pheromone-independent basal signaling

Unlike *sst2*∆ or *msg5*∆, *gpa1*∆ has increased basal signaling compared to wild type **(Figure 1D)**. We first attempted to rescue wild type signaling in *gpa1*∆ without correcting for this basal signaling difference. However, all open loop inductions of Opto-Gpa1 starting concurrently with pheromone induction failed in a prototypical way. Low constant expression of Opto-Gpa1 drove output down from its high basal state, eventually reaching the wild type level but only around two hours post induction (**Figure 4 trial 1**). Higher Opto-Gpa1 expression shortened this time, but at the cost of over attenuation of signaling due to over-production of Gpa1 (**Figure 4 trial 2**). These data suggested that basal expression of Gpa1 in cells prior to stimulation is required for proper signaling dynamics after α-factor induction.

**Figure 4.**
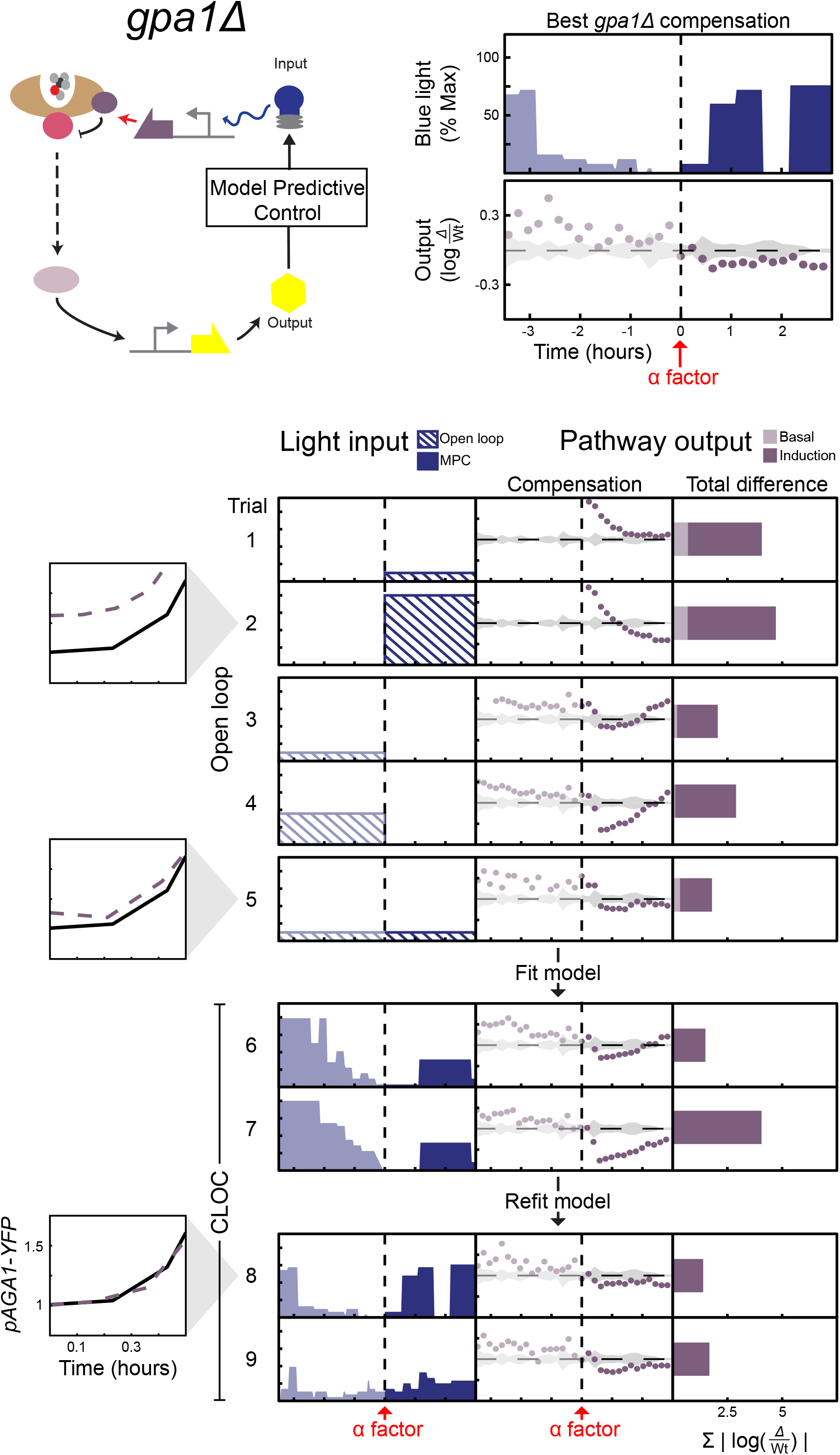
CLOC experiments indicate that the basal error for *gpa1*∆ needs to be corrected in order to restore post α-factor signaling dynamics. (a) For all trials, mutant strains are induced with 0.5nM α-factor at time 0 hours and *pAGA1-YFP* measured for time -3.5 to 0 hours (basal) and 0 to 3 hours (induction). Compensation of the mutant is carried out through optogenetic expression of the *GPA1*. Results of different trials are shown as rows in tables with three columns. The first column shows the light input as a function of time, the second column shows the resulting difference in signaling output as compared to wild type as a function of time, and the third column shows the sum of the differences shown in the second column. Basal difference (light purple) is the difference at the time of α-factor induction while the induction difference is the sum of all difference after α-factor induction (dark purple). Light inputs were given in either open loop (input is dashed), closed loop (input is solid color). Insets compare *pAGA1-YFP* output of the wild type (solid line) and *gpa1*∆ (dashed line) for the accompanying trial immediately post α-factor addition. Optogenetic input resulting in the best compensation along with its compensation error is shown on the top of the figure. This plot also shows the numerical values of the axes. The same scale and values are used for all panels. (See Figure S3, Figure S4, and Table S3)

Next, we asked whether constant open-loop expression of Opto-Gpa1 that precedes pheromone induction could rescue post-induction signaling dynamics in *gpa1*∆. We therefore induced Opto-Gpa1 for 3.5 hours before pathway activation by α-factor. After addition of α-factor, constant induction was either stopped (**Figure 4 trials 3–4**) or continued (**Figure 4 trial 5**). Light intensities strong enough to drive basal signaling in *gpa1*∆ to wild type levels resulted in over attenuation post pheromone induction, even in the absence of continued expression of Opto-Gpa1. While weaker constant induction of Opto-Gpa1 before α-factor addition avoided this over attenuation, it did not fully restore basal signaling in *gpa1*∆ (**Figure 4 trial 5 inset**) and if not continued post pheromone addition, also resulted in pathway over-activation at around 30 minutes after induction (**Figure 4 trial 5**).

We therefore hypothesized that dynamic compensation of *gpa1*∆ is necessary to fully rescue wild type basal signaling and immediate post induction dynamics without causing over attenuation. To perform CLOC of both basal signaling activity and induction dynamics, we parameterized a model using all the data generated by the constant light responses of *gpa1*∆ (**Figure S4A**). CLOC experiments using this initial model were able to satisfactorily rescue wild type basal signaling, generally through strong initial induction of Opto-Gpa1 that is subsequently ramped down prior to pheromone addition (**Figure 4 trials 6–7**). However, this induction strategy did not avoid the issue of over attenuation following α-factor addition seen in the constant Opto-Gpa1 experiments.

Using these data, we then performed a second round of model fitting for additional CLOC experiments. These experiments resulted in more tapered Opto-Gpa1 expression strategies that rescued basal signaling *gpa1*∆ (**Figure 4 trials 8 inset**) without causing severe over attenuation immediately post α-factor induction. The same light induction strategy generated similar results for *gpa1*∆ CLOC experiments conducted at a higher concentration of α-factor (**Figure S4C**). These data indicate that a carefully calibrated basal expression of Gpa1 is needed at the time of pheromone addition in order to start approaching post-induction wild-type dynamics (**Figure 4, compare trials 6 and 7 with trial 8**).

Furthermore, many quantitatively different Opto-Gpa1 inductions could rescue wild type *pAGA1-YFP* levels without extensive over-expression of Gpa1 at the time of pheromone induction. These strategies resulted in similar rescue of post-pheromone dynamics and showed only a weak dependence on post-pheromone Opto-Gpa1 expression (**Figure 4 compare trials 8 and 9, also compare trials 1 and 2 to trials 8 and 9**). Taken together, these data strongly suggests that negative feedback via transcription of *GPA1* plays a critical role in setting this level during basal signaling, and that this level is essential for pathway dynamics after pheromone induction.

## DISCUSSION

Feedback loops have mostly been studied in contexts where they generate dramatic *qualitative* behavior, such as the positive feedbacks that produce the sharp switching (Xiong and Ferrell, 2003) used for cellular differentiation or the negative feedbacks that produce oscillations (Pomerening, Kim and Ferrell, 2005) used in setting biological rhythms). It is widely accepted however, that hundreds if not thousands of feedback loops decorate a cell and that the large majority of these do not create cellular phenotypes but instead contribute *quantitatively* to them. Understanding these feedbacks requires knowing both their dynamics, when and how strongly they are active, and more importantly what happens when these dynamics are disrupted. Despite previous elegant attempts (Santos, Verveer and Bastiaens, 2007; Yu et al., 2008), systematic tools are scarce to efficiently generate hypotheses pertaining to these questions. The approach presented here takes a first step in this direction by combining two powerful ideas, genetic compensation and control theory. Adding a dynamic dimension to classical genetics by defining phenotype as time-dependent signaling, we ask control theory to determine how much and how often introduction of a deleted feedback component can rescue wild-type signaling dynamics in a mutant strain. We then use this time trace of the introduced compensatory feedback and its success or failure in rescuing wild type signaling to generate hypotheses about the dynamism, timescale of action, sensitivity, and other hallmarks of the endogenous feedback loop.

In the case of the mating pathway, we determined that the extensively studied feedback loop involving *SST2* whose dynamic induction in response to pheromone has largely been assumed to be essential for signaling (Dixit et al., 2014), appears to have no temporal requirement for generating wild type induction dynamics. Instead static expression of Sst2 is sufficient to generate wild type signaling. By contrast, our studies suggest that the feedback loop involving the phosphatase *MSG5* is exquisitely dynamic, pulsing repeatedly to fine-tune pathway dynamics. Our data also suggest that *GPA1* is involved in dynamic feedback, but one that is important mostly during basal signaling.

This operational definition of the three feedback loops could not be achieved by direct measurement of pheromone induced *GPA1*, *SST2*, and *MSG5* transcription dynamics alone. Typically, measuring an increase or decrease of a gene product in a given circumstance doesn’t causally imply its functional relevance to this circumstance. To make such a determination classically, one would observe the effect of a disruption to this gene and rescue this effect by compensation with the wild type allele. Our methodology provides a direct parallel to this in that it does not simply correlate a dynamic transcriptional profile with a signaling output, but tests the sufficiency of this profile to produce the signaling output. By doing so, we provide a measure of causality.

While the phenotype we adopt here is time-dependent signaling, the same framework is applicable to other quantitative phenotypes, such as noise regulation or dose response alignment (Yu et al., 2008). For example, Sst2 was postulated to be a noise regulator in the mating pathway (Dixit et al., 2014). By defining the phenotype to be rescued through induction of Opto-Sst2 as the difference in cell to cell variability of *sst2*∆ as compared to wild type, one can repeat CLOC modified appropriately to generate concrete quantitative hypotheses about the noise regulatory role of the *SST2* feedback loop. Similarly, while we use optogenetic transcriptional control, our methodology could easily accommodate a number of existing optogenetic tools that modulate a variety of post transcriptional processes including cAMP secondary signaling (Stierl et al., 2011) and subcellular localization of proteins (Yumerefendi et al., 2015), and thus be expanded to serve as a general tool for the study of different types of feedback regulation beyond transcription. This is further facilitated by the use of a simple empirical model to devise the “rescue” control sequence. This aspect defies biological intuition where it is often considered that mechanistic models are most useful, a reasonable statement when direct analysis of the model is intended. Since the hypothesis generating entity here is the rescue sequence itself, we limited the granularity of the model, using a generic representation that can produce such a rescuing sequence while still being generalizable to other systems and applications.

Overall, we believe CLOC to be a broadly applicable tool that can update the classical genetics toolbox with a method that facilitates the study of one of the most essential aspects of life—the ability to self-regulate through feedback.

## ACKNOWLEDGEMENTS

We thank Ignacio Zuleta for advice on the design and construction of the automated closed-loop control device and software and Eric Chow and the Center for Advanced Technology at UCSF for help with its fabrication. This work was funded in part by DARPA (HR0011–16–2–0045) and the Paul Allen Frontiers Group for H.E.-S and the National Institute of Health (R01GM71801) for H.M. Both H.E.-S and H.M. are Chan-Zuckerberg Investigators.

## AUTHOR CONTRIBUTIONS

All experiments were conceived of by P.H., H.M., and H.E.-S and performed by P.H. Data analysis was carried out by P.H. Manuscript was written and edited by P.H., H.M. and H.E.-S.

## STAR METHODS

### Contact for reagent and resource sharing

Further information and requests for resources and reagents should be directed to and will be fulfilled by the Lead Contact, Hana El-Samad. To request reagents please submit a form to UCSF at https://ita.ucsf.edu/researchers/mta.

### Experimental model and subject details

#### Saccharomyces Cerevisiae

##### Plasmid and strain construction

All plasmids and strains used in this study are listed in tables S1 and S2. Plasmids contain single integration constructs selectable using auxotrophic markers. They were linearized by digestion with Pme1 and transformed into yeast using standard lithium acetate protocols. Annotated sequences are available on Addgene. The cryptochrome optogenetic expression constructs were PCR amplified from Addgene plasmids 28244 and 28245 and modified using Gibson Assembly. The gene encoding the secreted protease *BAR1* (Manney, 1983) was knocked out in all reported strains to ensure that α-factor was not extracellularly degraded during experiments. This was done by amplifying a nourseothricin resistance cassette flanked on either side by 70 base pairs of homology to the genomic sequence immediately upstream and downstream of the *BAR1* gene and transforming this amplicon into yeast. *SST2*, *MSG5*, and *GPA1* deletions were similarly accomplished using the amplification of a kanamycin resistance cassette. To control for previously reported basal expression in the dark from the cryptochrome optogenetic system, feedback deficient mutants are compared to wild type strains transformed with a copy of the optogenetic expression system and the corresponding light inducible allele.

##### Media and growth conditions

Single colonies are picked from YPD (yeast extract, peptone, 2% glucose) agar plates into 50 mL of YPD media and grown overnight at 30 degrees Celsius to an optical density of 0.2. 8 mL of this culture is added to 22 mL of YPD in a 50 mL conical tube containing a magnetic stir bar and incubated on the yeast incubator array for 1 hour prior to the start of an experiment.

### Method details

#### Automated closed loop control device and software

The device hardware consists of custom fabricated, LED-enabled yeast incubator arrays coupled to a liquid handling module and a three-laser flow cytometer (LSR II, Becton Dickinson) (**Supplemental Information**). Each incubator array consists of four temperature controlled and magnetically stirred chambers designed for liquid culturing of yeast in 50 mL conical tubes (Falcon, Becton Dickinson). Temperature control is achieved using a thermocouple and PID regulated resistive heating. Stirring is achieved using magnets mounted to a rotor using a laser cut adapter to turn magnetic stir bars.

Chambers are light isolated and contain a heat sink on which three individual LEDs can be mounted. Custom fabricated caps allows for liquid to be moved in and out of the conical tubes using flexible tubing and standard one-piece fluidic fittings (PEEK No-Twist, VICI). Multiple incubator arrays can be attached to the liquid handling module, and we used 2 arrays for a total of 8 chambers in all experiments presented above. Liquid handling is accomplished using the two syringe pumps (Cavro XCalibur Pump, Tecan) of a High Throughput Sampler (Becton Dickinson) and a stream selector (Valco SD selector, VICI). The liquid handling module allows for the movement of yeast culture from an individual incubator to the cytometry for measurement or to a waste reservoir for culture dilution. Fresh media and α-factor can likewise be moved into individual incubators. A complete schematic of this fluidics setup is shown in the supplemental information.

The syringe pumps, stream selector, and LED driver are each serially addressable and are concurrently controlled during experiments by custom software (LabView, National Instruments) used to coordinate their individual actions. Additionally, this software processes the cytometry data generated during the experiment in real time. This data is then passed to a MatLab program that implements the model predictive control algorithm and dictates the appropriate optogenetic inputs to the LED driver. All model files required for fabrication of device parts are available upon request and a bill of materials is provided in the supplemental information.

#### Automated closed loop and open loop control measurements

The LabView software is used to specify for each experiment the number of culture samples, a sample frequency, a number of outgrowth measurements, and a dilution volume. At each measurement time point, culture is sampled iteratively from each incubator within the array. A sample consists of moving culture from an incubator into the cytometer for measurement, diluting by replacing culture with a volume of fresh media, and then clearing the fluidics lines by pushing an air bubble through the lines and into a waste reservoir. This clearing was sufficient to prevent mixing of subsequent samples (data not shown). For *sst2*∆ and *msg5*∆, we used a sampling frequency of 8 minutes and a dilution volume of 1mL. For *gpa1*∆ we used a sampling frequency of 12 minutes to accommodate the time required for the higher dilution volume of 2 mL needed to maintain exponential growth during the longer experiments. For each experiment, cultures are first measured for a specified number of outgrowth time points (4 for *sst2*∆ and *msg5*∆ and 18 for *gpa1*∆) in order to estimate basal pAGA1-YFP signaling. Before the first measurement after the outgrowth period, 1 mL of YPD containing α-factor is added to each incubator such that the final culture concentration is either 0.5 nM α-factor (**Figure 3–4**)or 1 nM α-factor (**Figure S4**) and the dilution media is replaced by YPD containing the appropriate concentration of α-factor. Light induction begins concurrently with either α-factor addition for experiments with *msg5*∆ and *sst2*∆ or the first culture measurement in the case of *gpa1*∆. In order to prevent phototoxicity during induction of the light inducible alleles, blue light is administered at the specified intensity during a 2 second pulse repeated every 2 minutes. Given the previously reported half-life of activation of the cryptochrome protein, these light pulse trains are sufficient to achieve constant activation. Two incubators within the array are always used as internal controls for α-factor concentration and receive no blue light during the duration of an experiment. A detailed description of protocol and sequential steps used by the automated device for sampling, cleaning, diluting, and induction including the settings for valve positions and syringe actions are included in the supplemental information.

### Quantification and statistical analysis

Cytometry measurements are processed by the device software in real time by discarding the first 750 events of a sample in addition to any events with no measured fluorescence. All fluorescence measurements (FITC) are then normalized to cell volume by dividing by side scatter (SSC) and *pAGA1-YFP* levels taken as the median FITC/SSC value of the population. These levels are reported as fold change over basal with the basal level estimated as the median of the measurements taken at the time of pheromone addition (0 hours). For a given time point, the difference in signaling between the wild type and mutant strain (compensation) is calculated as the log_2_ of the ratio of the mutant output to the wildtype output. The total difference is calculated by summing the absolute value of each time point difference that exceeds the standard deviation observed for biological repeats at that time (gray shaded region) in the wild type strain in the absence of light.

### Data and software availability

LabView code for control of automated yeast culturing, pheromone induction, cytometric measurement, and light delivery using the device described above is available upon request. A summary of the operations and algorithms used in this code is provided in the supplemental information.

KEY RESOURCES TABLE

KEY RESOURCES TABLE
| REAGENT or RESOURCE | SOURCE | IDENTIFIER |
| --- | --- | --- |
| Chemicals, Peptides, and Recombinant Proteins |  |  |
| Alpha-Factor Mating Pheromone | VWR Scientific | Cat #: H-2925.0001 |
| Experimental Models: Organisms/Strains |  |  |
| *S. cerevisiae* Strains, see Table S1 | This paper | N/A |
| Recombinant DNA |  |  |
| pGAL4BD-CRY2PHR | Addgene | Plasmid #: 28244 |
| pGal4AD-CIB1 | Addgene | Plasmid #: 28245 |
| Yeast integrating plasmids, see Table S2 | This paper | N/A |
| Software and Algorithms |  |  |
| LabView software for automated closed loop control device | This paper | N/A |
| Other |  |  |
| Automated closed loop control device | See Table S4 | See Table S4 |

